# Circular permutants of azurin exhibit molten globule intermediates not observed in WT azurin

**DOI:** 10.1101/2025.10.10.681544

**Authors:** Debanjana Das, Sri Rama Koti Ainavarapu

## Abstract

Molten globule (MG) states of proteins have been described as special types of intermediate states within the energy landscape diagram of proteins. In a cell, the MG state has functional significance compared to the native or unfolded states owing to its enhanced side-chain dynamics. In this study, we re-examined the acid-denatured state of the metalloprotein azurin and also investigated whether circular permutation (CP) of azurin can lead to modulation of the energy landscape to yield intermediate or MG states. CP is a protein engineering tool that allows the rearrangement of the secondary structural elements without perturbing the overall three-dimensional protein structure. We carried out comprehensive thermal, chemical, and pH denaturation studies to examine possible MG states in azurin WT. We additionally examined two CPs of azurin, whose energy landscapes in the metal-free and metal-bound forms have been previously characterized. One of the positions of CP is near the metal-binding loop (cpN42) and the other is within the active site of the protein (cpF114). We used thermal denaturation monitored by far-UV and near-UV CD, time-resolved fluorescence anisotropy of a buried Trp, and solvent accessibility of the same Trp residue to evaluate the presence of MG states in these proteins. WT azurin shows binding with ANS at a low pH due to ionic interaction with the fluorophore at this pH. We find that though WT azurin does not adopt an MG state, CP of azurin(s) exhibited an MG state. Our findings highlight the potential of CP to modulate protein energy landscapes and consequently the physiological functions of proteins.

## Introduction

Protein folding pathways can be studied by precise thermodynamic characterization of the folding intermediates of proteins.^1^ The energy landscape model explains the protein folding as an energetically downhill process from the denatured state to the global minima (native state) via intermediate states such as molten globule (MG) states.^2–4^ The MG state of a protein is a partially unfolded intermediate state with lost tertiary contacts but is still a compact form of the protein involving an intact secondary structure.^5–10^ MG states have been reported in a variety of proteins such as α-lactalbumin^11–18^, human serum albumin^19–21^, myoglobin^22,23^, barstar^24–26^, cytochrome c^27^, SUMO2^28^ (Small Ubiquitin-related MOdifier 2), azurin^29^, p53^30^ (tumor suppressor protein), β-lactamase^31^, carbonic anhydrase^32–35^, and many more^36–40^. The MG state in various proteins also has functional significance^6,41,42^ and is also capable of binding to other proteins^35,39^. The MG state of proteins has been shown to play key roles in protein folding as well as in protein aggregation and fibrillization.^43,44^ Interestingly, the MG state of human α-lactalbumin performs an important function in the apoptosis of tumor cells.^16–18^ Hence, the characterization of the MG state of proteins can help us to get insights into complex protein-folding and aggregation-associated problems.

Proteins often exhibit MG states in slightly denaturing conditions such as low pH^12,19,31,45^, mild denaturant^12,32^, high temperature^28,46–49^, or high pressure^50^. Conventional experiments utilized to examine the MG state of a protein are the binding of a hydrophobic dye molecule like 8-anilino-naphthalene-1-sulfonic acid (ANS) and structural characterization by far-UV and near-UV circular dichroism (CD). The absence of the tertiary structure with a retained secondary structure is the primary characteristic of the MG state. In MG states, the loss of the tight packing of side chains in the protein often results in exposure of the core hydrophobic region. The ANS molecule shows strong binding affinity with the MG state of the protein due to hydrophobic interactions with the exposed hydrophobic core of the protein.^51^ As a result, ANS gives enhanced and blue-shifted fluorescence as compared to that in polar solvents.^52^ On the other hand, native and denatured states of the protein have very poor binding affinity to ANS.^36^ A review by Hawe *et al.* describes the use of extrinsic fluorescent dyes (such as ANS) for the characterization of MG or aggregated states of proteins.^53^ Hence, an increase in fluorescence emission intensity of ANS acts as the marker of the MG state. While ANS is a useful and easily accessible probe for general MG characterization, maximal ANS fluorescence should not be the sole criterion for establishing the MG state of a protein. For example, previous studies on chymopapain showed maximal ANS fluorescence at pH 1.0 along with a huge loss (∼80%) of secondary structure, whereas at pH 1.5 it showed enhanced ANS fluorescence with secondary structure intact.^45^ Therefore in such cases, in the absence of any other characterization method, ANS might erroneously label the denatured proteins as MG states. The MG states of proteins have also been characterized by investigating Trp residues within proteins. The solvent exposure and mobility (using fluorescence anisotropy) of Trp in the native and MG states have been explored.^13–15,23–25,37,38,54,55^ Reports from the literature show that the solvent exposure of Trp increases on conversion from the native to MG state.^13,15,38,54,55^ Interestingly, several literature reports show that the mobilities of Trp in the native and MG states are similar.^13,23,37,38^

Azurin, a metalloprotein has been reported to exhibit a molten globule state at pH 2.6 by ANS binding.^29^ Azurin is a Cu-containing metalloprotein (Figure 1a) with anti-cancer properties ^56–58^ and is derived from bacteria such as *Pseudomonas aeruginosa*. The redox-active Cu-ion acts as an electron shuttling protein in the respiratory systems of such bacteria. The copper active site of the protein has five ligands (Gly45, His46, Cys112, His117, and Met121), amongst which the Cys112-Cu^2+^ is the strongest bond which also gives rise to an LMCT transition^59,60^ and the brilliant blue color of the protein. Azurin also has an intrinsic tryptophan (Trp48) residue in a highly buried environment^61^ and its (un)folding processes have been well-studied^62–65^. Azurin (both apo and holo) can bind to and stabilize the tumor suppressor protein p53.^56,66^ Hu *et al.* have recently solved the crystal structure of this azurin-p53 complex, establishing the residues involved in this interaction.^67^ They have further obtained several affinity-enhancing mutants to increase the apoptosis effect of azurin on cancer cells. In this regard, the energy landscape of the protein drug candidate azurin should be critically assessed. Any MG states or partially folded intermediates can affect the efficacy of the active form of the protein binding to p53. In a report, Sandberg *et al.* show that C112S and C112A metal-free azurins adopt MG states on refolding and the results are extrapolated to WT azurin.^29^

**Figure 1.**
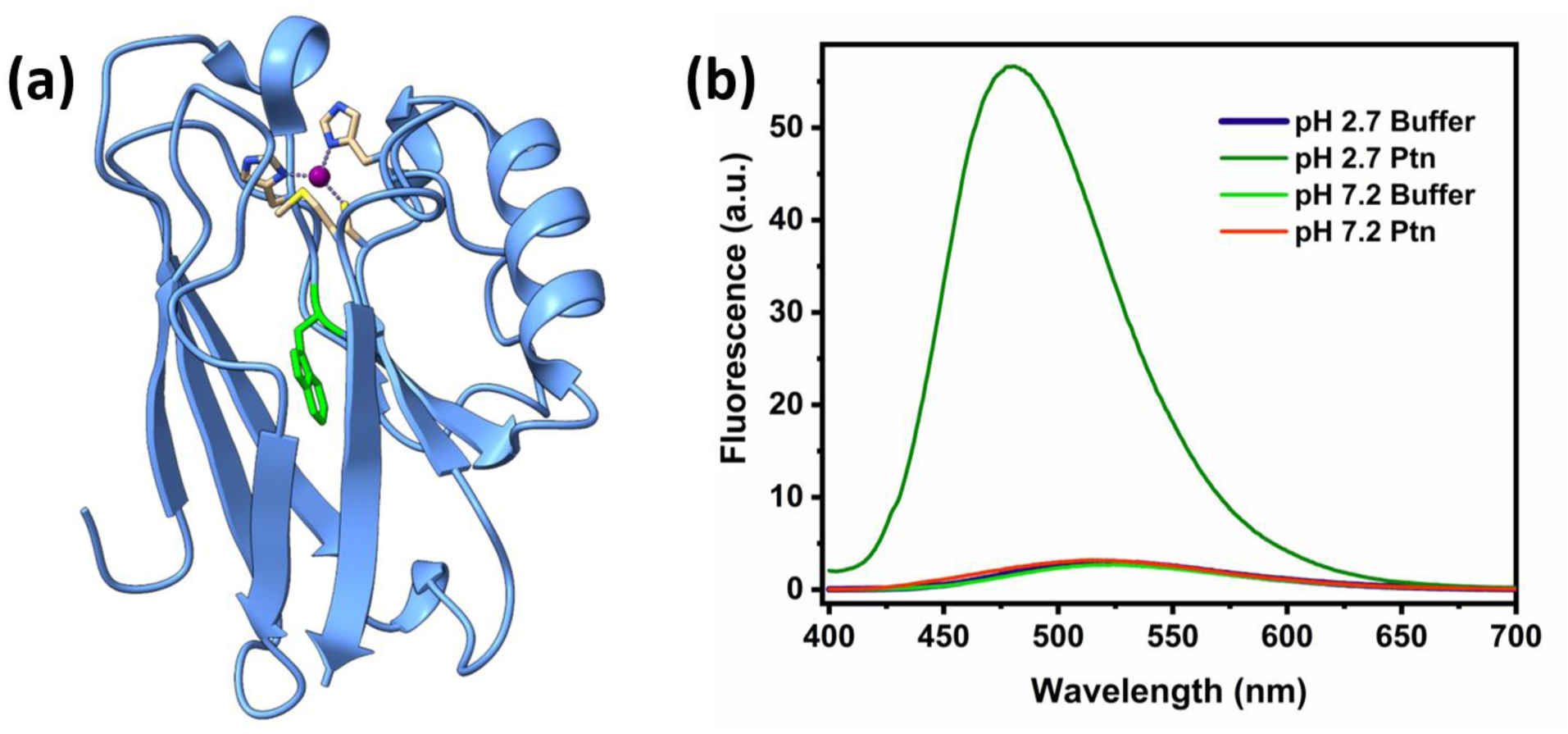
**(a)** Structure of Cu^2+^-azurin (PDB ID: 4AZU^94^) with the copper ion shown as a purple sphere and tryptophan side-chain in green. **(b)** Enhanced blue-shifted ANS fluorescence in Cu^2+^-azurin at low pH ([Protein] = 10 μM and [ANS] = 500 μM).

In the current study, we re-examine the pH-induced MG state of azurin WT described in the literature. This is relevant since previous experiments were performed on mutant azurins (C112S and C112A). Here we show, by systematic studies of the chemical and thermal denaturation of azurin, that WT azurin does not have MG states. The ANS-bound state of apo-azurin at low pH reported in literature^29^ is an unfolded state of the protein. The enhanced ANS fluorescence is due to ANS binding to positively charged residues in the protein at pH 2.6. Additionally, we investigate whether circular permutation (CP) can induce MG states in the azurin protein. CP rearranges the polypeptide sequence of a protein, however, keeping the tertiary fold of the protein the same. CP can introduce intermediates on the folding pathway.^68–70^ For our studies, we use cpF114 and cpN42 azurins, whose major structural characteristics are similar to the WT proteins.^71,72^ We have reported that the CP azurins exhibit intermediates in their folding landscapes.^73,74^ Zn^2+^-cpN42 exhibited an on-pathway intermediate^73^ and Zn^2+^-cpF114 exhibited off-pathway intermediates^74^ in the refolding pathway. Thus, we investigated the CPs of azurin that exhibited intermediates in the context of the introduction of MG states in the protein energy landscape. We utilized various experimental techniques such as ANS-binding, CD, time-resolved fluorescence anisotropy, and acrylamide quenching to investigate the presence of MG states in CP azurins and compare them with our studies on WT azurin. Zn^2+^-cpN42 azurin (has an on-pathway intermediate) showed increased mobility and solvent accessibility of a buried Trp residue in the protein compared to other azurins. Our findings highlight the potential of CP of proteins in modulating protein energy landscapes and consequently the physiological functions of proteins to understand protein energy landscapes and how modifications such as CP can facilitate the formation of MG states, thus enhancing our grasp of protein stability and folding pathways.

### Experimental section

#### Chemicals used and protein synthesis

Guanidine hydrochloride (GdnHCl) was procured from Sigma Aldrich. The concentration of the stock GdnHCl solutions was calculated using the refractive index method. ANS (8-anilinonaphthalene-1-sulfonic acid) was bought from Sigma Aldrich. Acrylamide was bought from Amresco Pvt Ltd. Azurin WT and cpF114 proteins were overexpressed and purified using the osmotic shock protocol.^73,75^ Azurin cpN42 was overexpressed and purified using sonication followed by Ni-NTA affinity chromatography as done previously.^73^ The pure proteins were obtained by using SEC (size-exclusion chromatography) on a Bio-Rad Biologic Duo-Flow SEC system with a superdex75 column (GE Healthcare). The buffer used for eluting holo-protein was 20 mM Tris at pH 7.4. Apo-protein was eluted in 20 mM Tris buffer containing 1 mM EDTA to avoid metal-bound protein. BSA (bovine serum albumin), lysozyme (from chicken egg white), and myoglobin (from equine heart) were procured from Sigma Aldrich. Ubiquitin and SUMO1 were overexpressed and purified from *E. coli* according to previous protocols.^76^ The purity of the protein samples was verified by MALDI-ToF (Matrix Assisted Laser Desorption Ionization-Time of Flight) mass spectrometry (Figure S1). All experiments were conducted at 25 ℃, except for thermal denaturation experiments.

#### Circular dichroism (CD)

The steady-state far-UV CD data were recorded on a Jasco J-1500 spectrometer in the range of 200-260 nm. Scan speed was kept at 50 nm/min with 3 accumulations. The near-UV CD was recorded in the range of 260-310 nm. Scan speed was kept at 50 nm/min with 5 accumulations. For temperature denaturation by CD, the range scanned was 25-95 ℃ and the temperature interval was set as 2 ℃. The protein concentration used for far-UV CD experiments was 20-25 μM and that for near-UV CD experiments was 150-200 μM. Secondary structure analysis was done using BeStSel analysis software.^77,78^

#### Steady-state and time-resolved fluorescence and time-resolved anisotropy

Steady-state fluorescence measurements were performed on a SPEX fluorolog T-format fluorimeter (Horiba Jobin Yvon). Tryptophans of the proteins were excited at 295 nm and fluorescence was collected from 300-450 nm. Samples with different GdnHCl concentrations were prepared as explained previously.^73^ The concentration of protein in samples was 10 μM and the concentration of ANS was 500 μM. ANS was incubated with the protein for a minimum of 15 minutes. ANS was excited at 385 nm and fluorescence emission was collected from 400-700 nm. For acrylamide quenching experiments, the 4 (M) acrylamide stock solution was made in water. Titrations with acrylamide were made by adding 1 μL of the stock acrylamide solution to a 400 μL protein solution to introduce 10 mM acrylamide to the sample. F_o_/F (at 307 nm and 355 nm for NATA) vs [Acrylamide] was plotted for the different protein samples and fitted to Stern-Volmer equation (1)^28^. Here *F*_*o*_ is the initial fluorescence without the quencher, *K*_*sv*_ is the Stern-Volmer quenching constant, and [*Q*] is the quencher concentration. In Eq. (2), *K*_*sv*_ can be represented as *K*_*Q*_τ_0_, where *K*_*Q*_ is the bimolecular quenching constant and τ_0_ is the intensity averaged lifetime in the absence of the quencher.

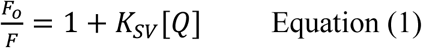

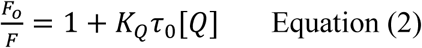

Time-resolved fluorescence and anisotropy for tryptophan were done by time-correlated single photon counting (TCSPC) as explained previously.^73^ The samples were excited at 295 nm and emission was collected at 340 nm. The instrument response function (IRF) had a full-width half-maximum of 80-100 ps. Mean fluorescence lifetimes have been estimated as described previously^79^, as 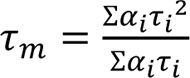. The time-resolved anisotropy can be written as a function of the initial anisotropy (r_0_), the amplitudes of the local and global motion of the fluorophore (α_1_ and α_2_), and the rotational correlation times (τ_R1_ and τ_R2_) as equation (3), where α_1_ + α_2_ = 1. Here we consider that the first term arises due to the segmental motion of Trp (τ_R1_ is for local motion) and the second term arises due to Trp rotation along with the macromolecule (τ_R2_ is for global motion).

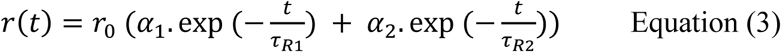

Trp can be considered to be rotating in the protein inside a cone of semi-angle α. The semi-angle α can be expressed in terms of the coefficients of the above equation (3) as per the following equation (4).^23,80,81^ The semi-angles of rotation of Trp in the protein cones of WT and CPs are calculated and given in Table S4.

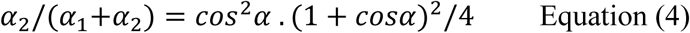

#### Stopped flow ANS fluorescence

The kinetics of ANS fluorescence was measured using a Biologic SFM300 stopped-flow fluorimeter as described in reference^73^. Protein, buffer, and ANS were excited at 385 nm and fluorescence was monitored by total integrated fluorescence above 455 nm. The concentration of protein in the cuvette after mixing was ∼10 μM. Concentration of ANS was kept at 500 μM in buffer, GdnHCl, and protein solutions.

## Results

### Investigation of MG states in WT azurin

Partially denatured or acid-forms of some proteins can exhibit MG states.^12^ Sandberg *et al*. have reported that azurin exhibits an MG state at pH 2.6, as probed by ANS binding.^29^ They have also observed ANS binding to mutant forms of apo-azurin (C112A and C112S) during refolding at pH 7.4.^29^ The physiological form of azurin is the copper-bound state. We find that indeed Cu^2+^-azurin shows high ANS binding (enhanced blue-shifted fluorescence) at pH 2.6, which is negligible at pH 7.4 (Figure 1b). Subsequently, we ask whether azurin exhibits reduced tertiary structure and/or ANS binding in mild chemically denaturing conditions, at pH 7.4. We thus subject Cu^2+^-azurin to a range of denaturant concentrations to shift the equilibrium from the folded state. Far-UV (monitoring secondary structure) and near-UV CD (monitoring tertiary structure) show that there is very little change by adding denaturant up to 2.0 M GdnHCl (Figures 2a and 2b). The c_m_ of Cu^2+^-azurin is 2.4 M, indicating that the protein will not be unfolded at 2.0 M GdnHCl. ANS fluorescence is not changed in the native state as compared to the partially unfolded state (Figure 2c). We also monitor ANS fluorescence change with the unfolding of Cu^2+^-azurin and observe that ANS fluorescence does not change with time (Figure 2d). Similar to the results for Cu^2+^-azurin, Zn^2+^-azurin also exhibits no ANS binding in mild-denaturing conditions though ANS binding occurs at low pH (Figure S2). These experiments hint at the absence of MG states in Cu^2+^-and Zn^2+^-azurins by GdnHCl denaturation.

**Figure 2.**
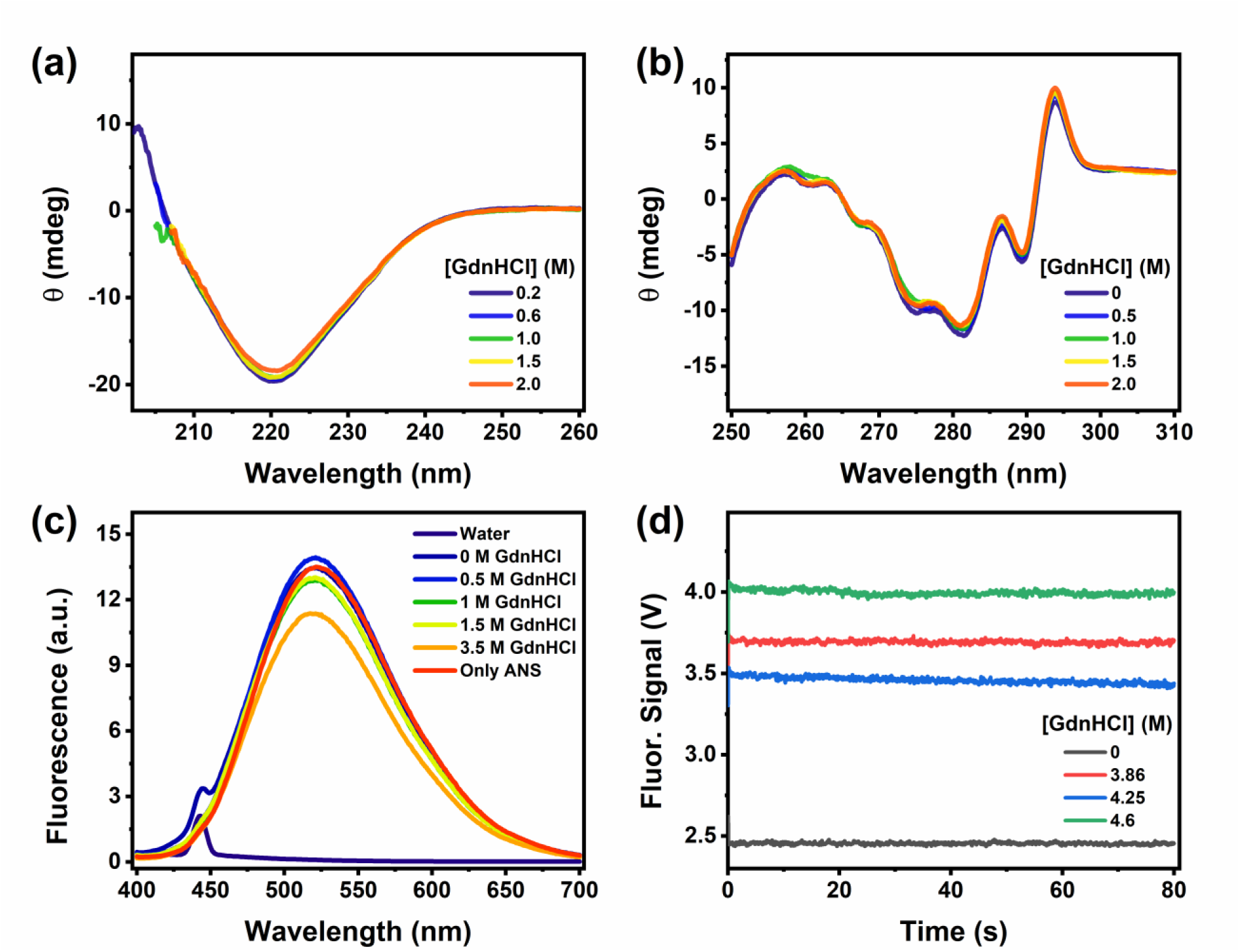
Chemical denaturation of Cu^2+^-azurin WT probed by **(a)** far-UV CD, **(b)** near-UV CD, and **(c)** steady-state ANS fluorescence shows no change in protein structure or ANS fluorescence. **(d)** Kinetic of ANS fluorescence shows no change in ANS fluorescence on the unfolding of Cu^2+^-azurin WT.

Since MG states have been observed at elevated temperatures ^28,46–49^, we chose to monitor the thermal melting of azurin with far-UV and near-UV CD. The change in CD signal was monitored at wavelengths 220 nm and 294 nm for far-UV and near-UV CD, respectively. The thermal denaturation profiles of Cu^2+^-bound, Zn^2+^-bound, and apo-azurin at pH 7.4 are given in Figures S3 and S4, respectively. Interestingly, the near-UV CD signal concomitantly changes with the far-UV CD signal, suggesting the absence of MG states in the samples. Therefore, the MG states are not observed upon heating the apo- or holo-azurins. We have systematically studied azurin in the acid-denatured form (pH 2.6) as per the literature report of azurin’s MG state at low pH.^29^ In our study, we examine the pH-induced MG state of azurin described in the literature further and ask whether this form of azurin exhibits reduced tertiary structure and/or ANS binding. The thermal denaturation profiles of Cu^2+^-azurin and Zn^2+^-azurin at pH 2.6 are given in Figures S3 and S4, respectively. For both these proteins, we observe that the melting point is significantly less (by >30℃) at the lower pH. The near-UV CD signal is not lost before the far-UV CD signal, suggesting that the near-UV CD is retained along with the far-UV CD.

We ask whether this acid form of azurin exhibits reduced tertiary structure and/or ANS binding. We find that at pH 2.6, Cu^2+^-azurin shows enhanced ANS fluorescence intensity with a blue shift of the emission wavelength to 480 nm (Figure 3a). Zn^2+^-azurin shows high ANS binding (enhanced blue-shifted fluorescence) at pH 2.6 and negligible ANS binding at pH 7.4 (Figure 3a). However, both the secondary structure and tertiary structure of Cu^2+^-azurin are retained (Figures 3b and 3c). Despite ANS binding, we observe that the tertiary structures of Cu^2+^-azurin and Zn^2+^-azurin are retained (Figure 3c). Contrastingly, apo-azurin shows a significant loss of far-UV and near-UV CD at pH 2.6, though there is a high extent of ANS binding (Figure 3). Additionally, the tryptophan fluorescence at ∼360 nm indicates that apo-azurin is unfolded at pH 2.6 (Figure 3d).

**Figure 3.**
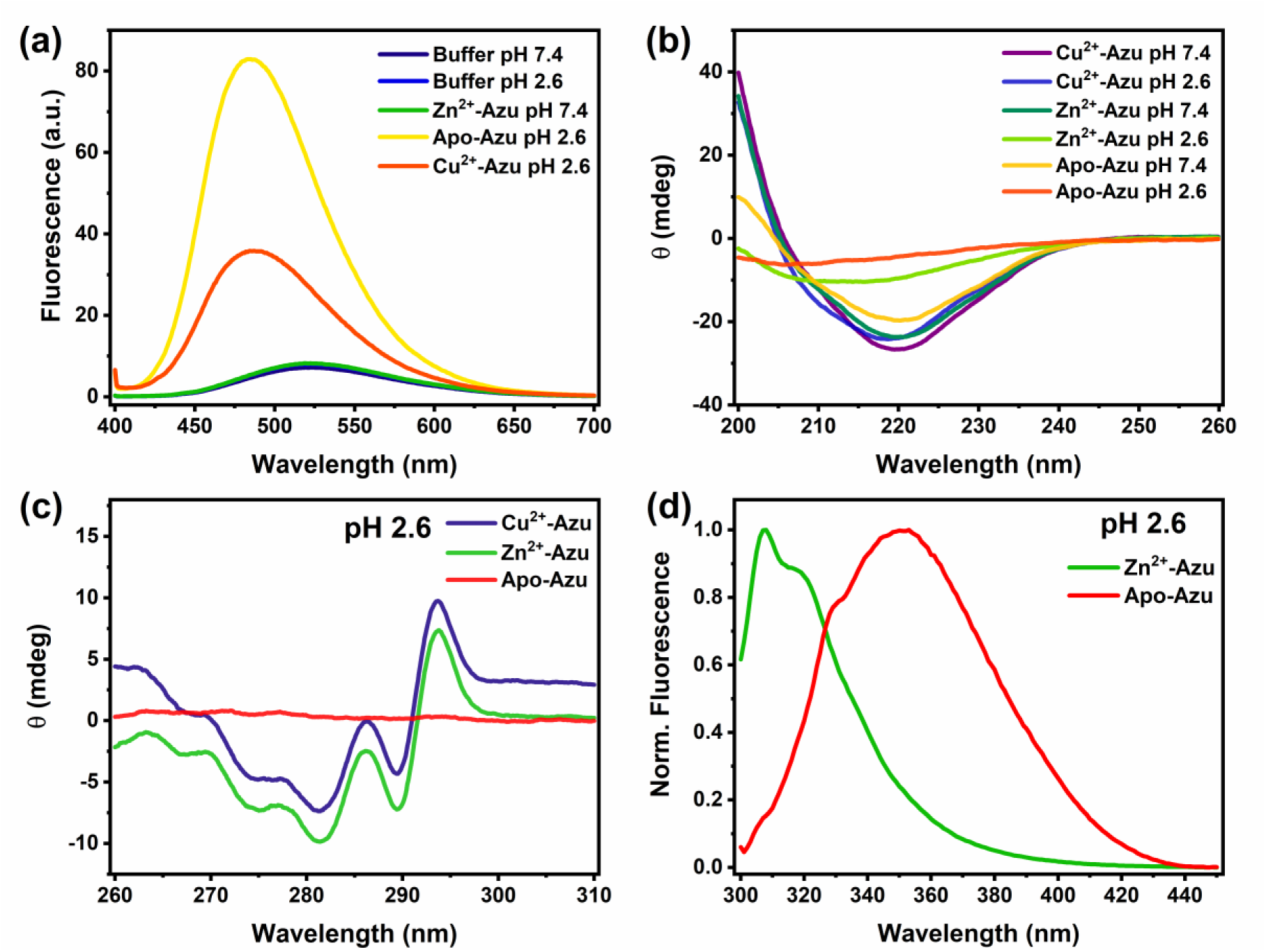
**(a)** ANS fluorescence on incubating with Zn^2+^-azurin WT (pH 7.4), and Cu^2+^- and apo-azurin WT (pH 2.6) shows that the proteins bind to ANS at the lower pH. **(b)** Far-UV CD of Cu^2+^-, Zn^2+^-, and apo-azurin WT at pH 2.6 and pH 7.4. (Protein concentrations for (b) are 20-25 μM). **(c)** Near-UV CD of Cu^2+^-, Zn^2+^-, and apo-azurin WT at pH 2.6. **(d)** Tryptophan fluorescence of Zn^2+^- and apo-azurin WT at pH 2.6. Far-UV CD, near-UV CD, and fluorescence show that apo-azurin is unfolded at pH 2.6.

Thus, we concluded that the ANS-bound state of apo-azurin at low pH reported in the literature ^29^ is an unfolded state of the protein.

### ANS binding to proteins at low pH

In the previous section, we observed that ANS binds to azurin at pH 2.6 and exhibits enhanced fluorescence. However, at physiological pH of 7.4, ANS shows quenched fluorescence and does not bind to azurin. We have performed a pH titration to see the pH range over which ANS gives high fluorescence with azurin as shown in Figure S5a. The resulting plot of fluorescence and pH (Figure S5b) shows that at lower pH values, the fluorophore binds to the protein and shows enhanced fluorescence. In fact, in literature, it has been observed that some proteins like bovine serum albumin (BSA), lysozyme, papain, and protease omega exhibit ANS binding at low pH.^82^ This is attributed to electrostatics and ion pair formation of the sulphonate group of ANS with positively charged amino acids in the proteins.^82^ These results indicate that at lower solution pH, the major contribution to the ANS fluorescence is due to the ion-pair interaction between the protein and ANS. To assess the generality of the enhanced ANS fluorescence at lower pH, we conducted ANS fluorescence studies at both pH 2.6 and 7.4 across a variety of proteins. For this purpose, we chose standard proteins such as BSA, lysozyme, myoglobin, and small globular proteins such as ubiquitin and SUMO1 (small ubiquitin-like modifier 1). In all the proteins studied, the ANS binding was shown to be enhanced at the lower pH (Figure 4). These results strongly suggest that ANS binding-induced enhancement in acidic pH is not necessarily indicative of MG states.

**Figure 4.**
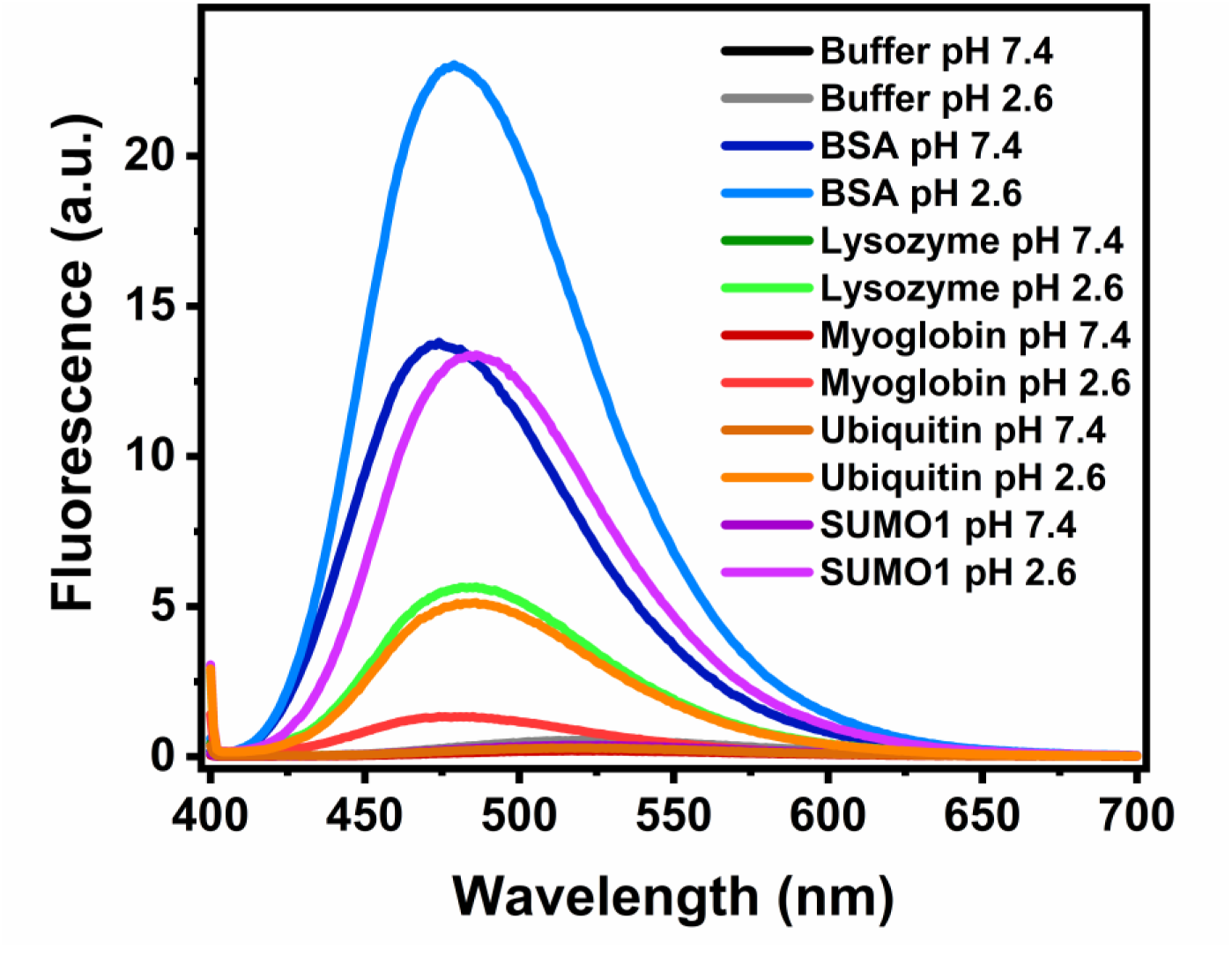
ANS fluorescence in BSA (bovine serum albumin), lysozyme, myoglobin, ubiquitin, SUMO1 (small ubiquitin-like modifier), and buffer at pH 7.4 and 2.6 ([Protein] = 10 μM and [ANS] = 500 μM) shows that ANS binds more to the various proteins at the acidic pH.

### Investigation of MG states in CP azurins

To explore how the connectivity of a protein chain influences ANS binding in response to changes in pH and chemical denaturation, we investigated ANS binding to circular permutants of azurin. Circular permutation (CP) consists of a change in the order of the secondary structure, essentially keeping the amino acid composition of the protein the same. CP was carried out at residue Asn42^72^ near the active site in the wild-type (WT) azurin protein and the protein was named cpN42-azurin^73^. Another CP was carried out within the active site at Phe114^71^ and was named cpF114-azurin^74^. Both the cpN42 and cpF114 azurins have Cu^2+^-bound structures similar to that of Cu^2+^-WT azurin.^71,83^ There are slight changes to the secondary coordination sphere around the copper. However, the overall structural features, including the hydrophobic core, are retained in the CP proteins. In the Cu^2+^-cpN42 azurin, the chemically denatured protein showed minimum ANS binding (Figure S6a), similar to WT azurin. However, the protein at a low pH shows high ANS binding as compared to that at physiological pH (Figure S6b). The enhancement of ANS fluorescence from pH 7.2 to pH 2.7 shown in cpN42 was larger than the corresponding enhancement for the WT-azurin protein. Given that the amino acid composition of WT and cpN42 was quite similar, this was presumably due to the presence of an extra His_6_ tag at the protein N-terminus of cpN42 azurin (Table S1). The His_6_ tag at the acidic pH would be positively charged and bind to negatively charged ANS molecules. Thus, we conclude that to determine that a particular protein has a molten globule state, ANS binding is not sufficient but one also should compare the far-UV and near-UV CD spectral changes to monitor the interior environment of the protein.

We have earlier reported that cpN42 when zinc-bound, has an on-pathway intermediate, not originally present in WT azurin.^73^ Also, we have recently demonstrated that apo- and Zn^2+^-cpF114 are highly destabilized CPs of azurin compared to azurin WT.^74^ Zn^2+^-cpF114 exhibited an off-pathway refolding intermediate. Both cpN42 and cpF114 azurins showed lower thermodynamic stability compared to WT azurin. Thus, we investigated the occurrence of MG intermediates in apo- and Zn^2+^-cpF114 and Zn^2+^-cpN42 by ANS-binding, CD (far-UV and near-UV), time-resolved anisotropy, and acrylamide quenching. The secondary structure analysis of apo-WT, apo-cpN42, apo-cpF114, Zn^2+^-cpN42, and Zn^2+^-cpF114 is given in Table S2. ANS-binding monitored with a change in temperature showed that apo-cpF114 showed a steady increase and then a decrease in ANS fluorescence with temperature (Figures 5a and 5b). This suggests that apo-cpF114 may have an MG state that prefers to bind to ANS. However, apo-azurin WT and Zn^2+^-cpF114 do not show the same behavior on ANS-binding monitored with increasing temperature (Figure S7). On the other hand, Zn^2+^-cpN42 shows that the ANS fluorescence decreases with temperature (Figures 5c and 5d). This may indicate that the Zn^2+^-cpN42 at room temperature is in the molten globule state. Chemical denaturation of Zn^2+^-cpN42 by GdnHCl also shows that at 0 M of the denaturant, the ANS binding or ANS fluorescence is maximum (Figure S8). However, Zn^2+^-cpN42 had a similar thermal melting curve monitored by far-UV and near-UV CD (Figure S9). Apo-cpF114 on adding Zn^2+^-ion shows enhanced far-UV and near-UV CD signal. Near-UV CD shows considerably more enhancement in signal than far-UV CD on adding Zn^2+^ to apo-cpF114 (Figure S10). This indicates that apo-cpF114 had significantly more loss of the tertiary structure as compared to the secondary structure, characteristic of an MG state.

**Figure 5.**
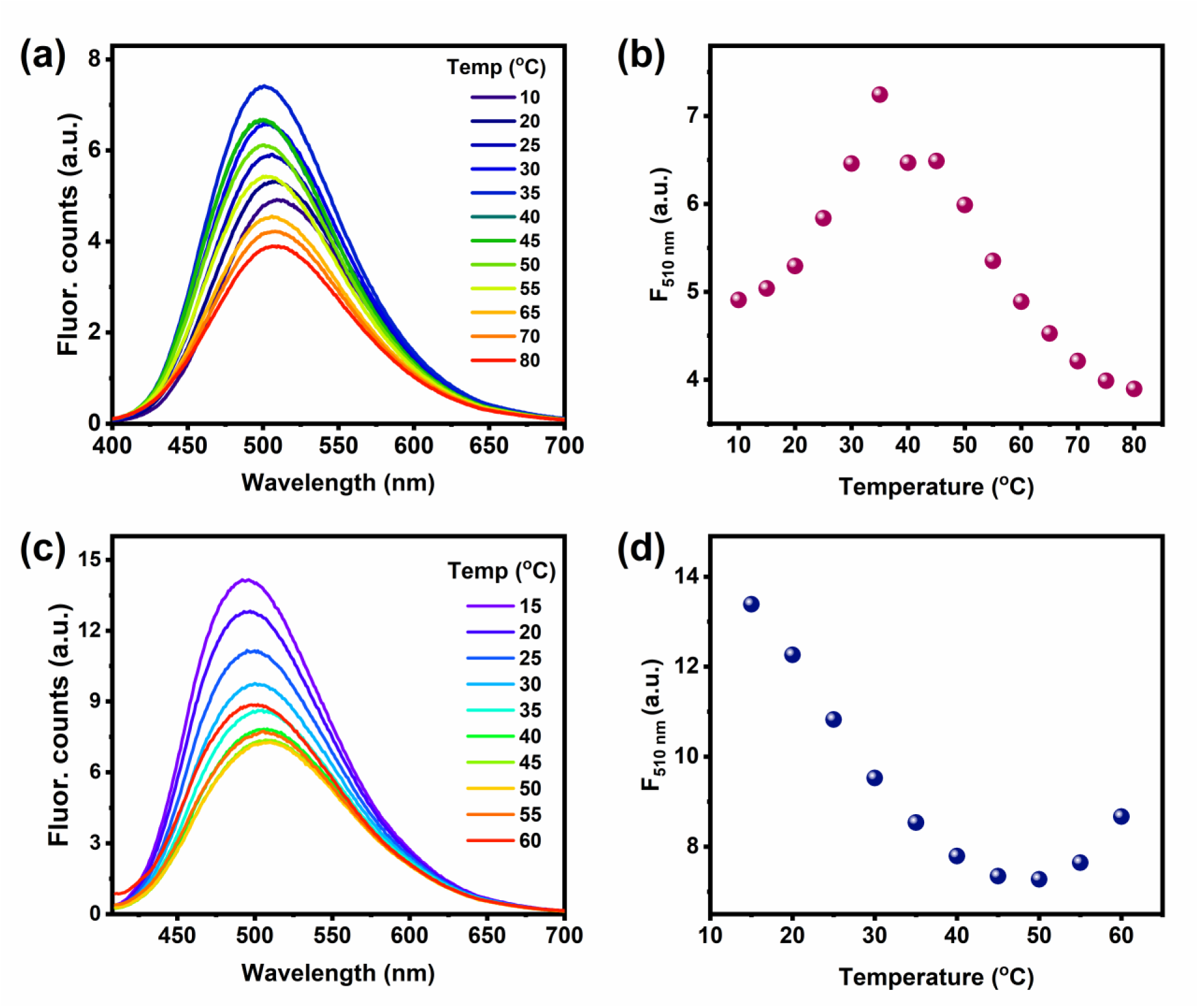
**(a)** ANS fluorescence response of apo-cpF114 azurin with temperature change. **(b)** Fluorescence change of (a) at 510 nm with temperature. ([Protein] = 30 μM and [ANS] = 500 μM). **(c)** ANS fluorescence response of Zn^2+^-cpN42 azurin with temperature change. **(d)** Fluorescence change of (c) at 510 nm with temperature. ([Protein] = 30 μM and [ANS] = 500 μM)

**Figure 6.**
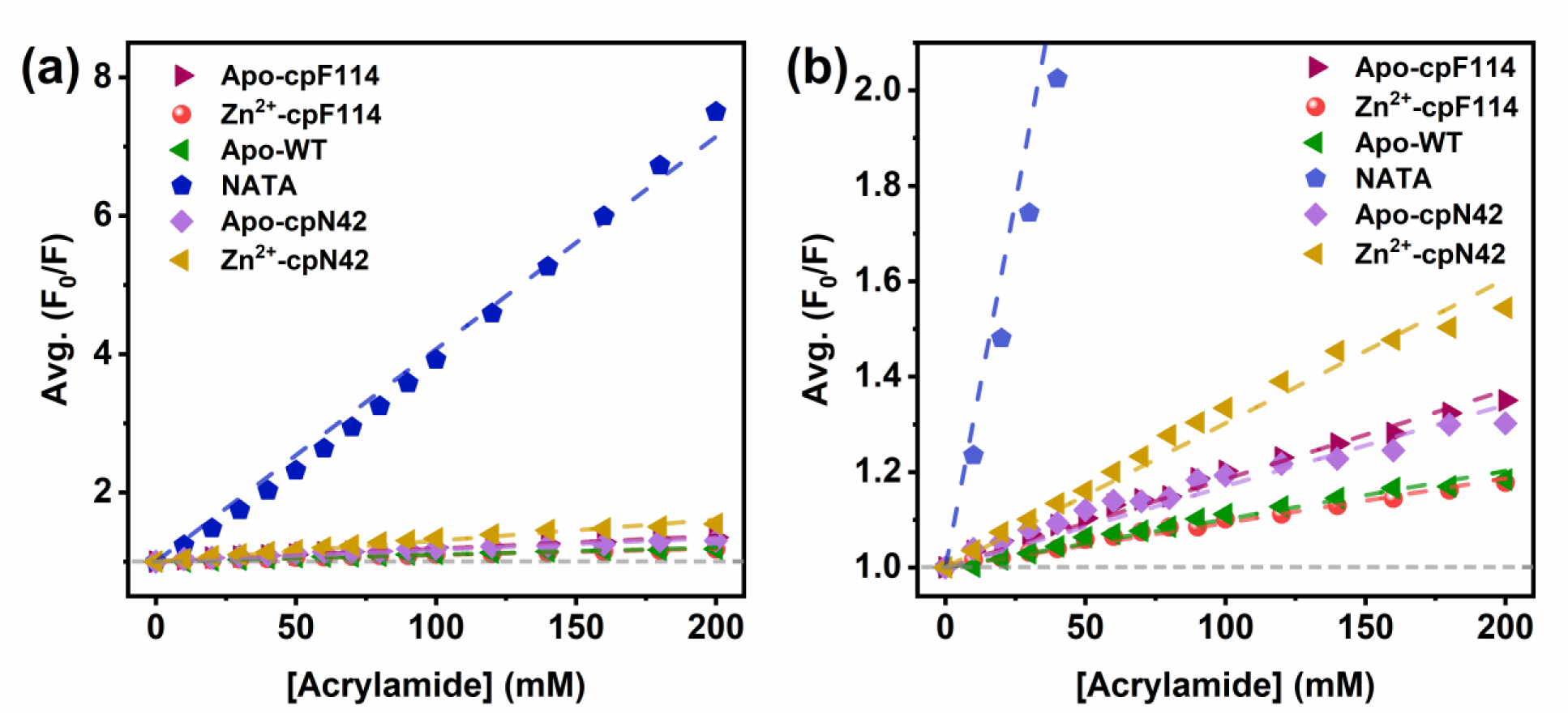
Stern-Volmer plots of acrylamide quenching data for apo-cpF114, Zn^2+^-cpF114, apo-WT, Zn^2+^-cpN42 azurins, and NATA show the relative accessibility of Trp in the various azurin mutants.

The MG states of proteins have been investigated by monitoring the Trp residues within the protein. The mobility of Trp and solvent exposure in the native and MG states has been previously explored.^13–15,23–25,37,38,54,55^ To further confirm the observations from ANS binding and CD, the lifetime and mobility (using fluorescence anisotropy) of the tryptophans in the WT and CP azurins were measured by time-resolved fluorescence (Figures S11 and S12). The components of rotational correlation times of Trp and their corresponding amplitudes are given in Table 1. The components of lifetimes are given in Table S3. Time-resolved anisotropies were fitted to two exponentials, denoting the global motion of tumbling the proteins and the local motions of Trp. The global motion has a higher rotational correlation time (4-6 ns) and the local motion has a lower rotational correlation time (≤ 0.1 ns). The amplitude of the local component of Trp motion in apo-cpF114 (0.65±0.03) and Zn^2+^-cpF114 (0.64) was higher as compared to that in apo-WT (0.54±0.07). Trp can be considered to be rotating in the protein inside a cone of semi-angle α. The semi-angle α can be expressed in terms of the amplitude coefficients. The cone of semiangle of rotation of Trp for apo-WT, apo-cpF114, and Zn^2+^-cpF114 were estimated to be 40°, 46°, and 45°, respectively (Table S4). This suggested that the mobility of the Trp for apo-cpF114 and Zn^2+^-cpF114 are the same. Several reports in the literature suggest that the mobilities of Trp in the native and MG states are the same.^13,23,37,38^ The amplitudes of Trp’s local component in apo-cpN42 and Zn^2+^-cpN42 are 0.48±0.04 and 0.81±0.02 respectively. Accordingly, the calculated semi-angles of rotation of Trp for Zn^2+^-cpN42, apo-cpN42, and apo-WT are 56°, 37°, and 40° (Table S4). Zn^2+^-cpN42 had the highest amplitude of the local component of Trp motion, indicating that at room temperature, Zn^2+^-cpN42 has a highly flexible Trp.

**Table 1.** Amplitudes and rotation correlation times obtained from the time-resolved anisotropy data of azurin-WT, azurin-cpF114, and azurin-cpN42 collected at 340 nm wavelength.

| **Protein** | **$\alpha_1$** | **$\tau_{R1}$ (ns)** | **$\alpha_2$** | **$\tau_{R2}$ (ns)** |
| --- | --- | --- | --- | --- |
| Apo-WT | $0.54 \pm 0.07$ | $0.07 \pm 0.01$ | $0.46 \pm 0.07$ | $4.59 \pm 0.90$ |
| Apo-cpF114 | $0.65 \pm 0.03$ | $0.04 \pm 0.01$ | $0.35 \pm 0.03$ | $4.18 \pm 0.53$ |
| Zn<sup>2+</sup>-cpF114 | 0.64 | 0.10 | 0.36 | 5.16 |
| Apo-cpN42 | $0.48 \pm 0.04$ | $0.10 \pm 0.02$ | $0.52 \pm 0.04$ | $5.66 \pm 0.80$ |
| Zn<sup>2+</sup>-cpN42 | $0.81 \pm 0.02$ | $0.06 \pm 0.01$ | $0.19 \pm 0.02$ | $4.80 \pm 1.17$ |

Data of apo-WT and apo-cpN42 are taken from reference<sup>73</sup> and data of apo-cpF114 and Zn<sup>2+</sup>-cpF114 are taken from reference<sup>74</sup>.

In an MG state, it is expected that the solvent exposure of the core residues increases. This can be verified by quenching of Trp fluorescence with a small quencher such as acrylamide, KI, or TCE (trichloro ethanol). These quenchers can access relatively solvent-exposed Trp residues. Reports from the literature show that the solvent exposure of Trp increases on conversion from the native to MG state.^13,15,38,54,55^ Only in rare cases is the opposite effect observed (W118 of human α-lactalbumin).^13^ We have carried out acrylamide quenching of the buried fluorophore (Trp) to find out the relative solvent exposure of the Trp in the different proteins. The Stern Volmer constant (K_SV_) calculated from the acrylamide quenching experiments are given in Table 2. A free fluorophore, NATA, shows a high K_SV_ value (30.7 M^-1^) due to high exposure to the quencher. The obtained values of K_SV_ are in the range observed in the literature. Apo-cpF114 has a much higher K_SV_ value (∼1.9 M^-1^) than Zn^2+^-cpF114 (K_SV_ value of ∼0.9 M^-1^) and apo-WT (K_SV_ value of ∼1.0 M^-1^), which confirms that the Trp in apo-cpF114 has a much higher solvent exposure. The Zn^2+^-cpN42 has yet higher solvent exposure of the Trp residue, as seen by the higher K_SV_ value (∼3.0 M^-1^). Apo-cpN42 has a lower solvent exposure of the Trp residue than Zn^2+^-cpN42 (K_SV_ value of ∼1.7 M^-1^). We have also used the excited state lifetime (τ_m_) of the fluorophore to find the bimolecular quenching constants (Table 2). The K_Q_ of apo-cpF114 (4.93*10^8^ M^-1^s^-1^) and apo-cpN42 (4.15*10^8^ M^-1^s^-1^) is much higher than Zn^2+^-cpF114 (2.42*10^8^ M^-1^s^-1^) or apo-WT (2.52*10^8^ M^-1^s^-1^), and that of Zn^2+^-cpN42 (7.46*10^8^ M^-1^s^-1^) is the highest. The K_Q_ value for apo-cpF114 is 100% higher than that of apo-WT. The K_Q_ value for Zn^2+^-cpN42 is 200% higher than that of apo-WT. These results in combination show that the proteins apo-cpF114 and Zn^2+^-cpN42 have higher solvent exposure of the core than the WT azurin.

**Table 2.** Stern-Volmer quenching constants (K_SV_) and bimolecular quenching constants (K_Q_) derived from acrylamide quenching experiments.

| **Protein** | **$K_{SV}$ (M<sup>-1</sup>)** | **$\tau_0$ (ns)** | **$K_Q$ (M<sup>-1</sup> s<sup>-1</sup>) \*10<sup>8</sup>** |
| --- | --- | --- | --- |
| Apo-WT | 1.01 | 4.00 | 2.52 |
| Apo-cpF114 | 1.86 | 3.77 | 4.93 |
| Apo-cpN42 | 1.71 | 4.12 | 4.15 |
| Zn<sup>2+</sup>-cpF114 | 0.94 | 3.88 | 2.42 |
| Zn<sup>2+</sup>-cpN42 | 3.02 | 4.05 | 7.46 |
| NATA | 30.70 | 2.85 | 107.72 |

## Discussion

### Importance of MGs in protein function and their characterization

MG states are thought to play important roles in several physiological processes^9,84,85^ due to their increased side-chain mobility.^42^ α-lactalbumin has been observed to exhibit MG states on the application of various conditions such as mild denaturant, pH, and temperature. Even in physiological conditions, complexed α-lactalbumin with oleic acid forms an MG and can induce apoptosis in a variety of tumor cells.^16^ Multiple engineered proteins are also reported exhibiting MG states.^85^

Azurin is a metalloprotein, well-known for its anti-cancer activity^56–58,67^, unique absorption^59,60^ and fluorescence properties^61^. The energy landscape of azurin has also been elucidated.^62–65^ In light of azurin as a promising cancer drug, the energy landscape of azurin and possible intermediate states are important. Mutant azurins (C112S and C112A) in the metal-free form showed MG states on refolding and the results were extrapolated to WT azurin.^29^ ANS binding kinetics and ANS fluorescence (at pH 2.6) support these claims. In our studies, we observe that azurin along with many standard proteins exhibit increased ANS fluorescence at lower pH. Thus, we advise exercising caution while analyzing ANS binding to proteins at low pH. As per previous studies, the ionic interaction of ANS with positively charged amino acids causes an increase in ANS fluorescence that is not necessarily due to MG formation.^82^ Indeed, there are reports where maximal ANS fluorescence at low pH does not correspond to the presence of MG states.^45^ Therefore, ANS fluorescence alone is not sufficient to comment on the presence of MG states. The use of multiple techniques such as far-UV CD, near-UV CD, anisotropy, and Trp fluorescence quenching by acrylamide provide a more holistic approach to examining MG states.

### CP as an engineering tool for introducing MG states in protein energy landscapes

Previously, CP has been reported to modulate the energy landscape of proteins by changing the (un)folding rates and/or by introducing intermediates.^68–70,73,74,86^ CP can also affect transient partial unfolding in proteins and alter the rate of native-state proteolysis. CP and WT DHFR proteins have differing tendencies to undergo transient partial unfolding.^83^ There has been literature evidence of CP of S6 that does not introduce MG in the protein.^87^ In other proteins such as dihydrofolate reductase (DHFR), CP introduces MG states in the CP proteins.^88,89^ However, ligand binding returned the CP protein to the native state with a tertiary structure. Investigating MG in azurin CPs underscores the importance of protein sequence and termini locations, especially in metalloproteins. We find that both Zn^2+^-cpN42 and apo-cpF114 exhibit higher Trp mobility and solvent accessibility as compared to WT azurin. These results can be further verified by probing the side chain dynamics of the residues in apo-cpF114 and Zn^2+^-cpN42 by NMR studies. Here, the residues responsible for the formation of the molten globule can be derived.

Contact order is a parameter used to estimate the extent of long-distance contacts required to form folded proteins. The relative contact orders (RCOs) for WT, cpN42, and cpF114 azurins (calculated from the formula in reference^90^) are 16.1%, 14.0%, and 17.1%, respectively. There does not seem to be an apparent relation of the CO with the propensity for MG state formation in azurin. Comparing CP proteins to WT proteins provides insights into evolutionary adaptations and the importance of MG states in protein function and evolution. Designing robust proteins for genetic engineering^91^, optogenetics^92^, or catalysis^93^ often involves circular permutation. Investigating MGs in CP proteins not only provides insights into the delicate balance between protein stability and flexibility but also further enhances and serves as a novel platform for making robust protein structures for biomedical and biotechnological applications.

## Conclusions

The MG state of a protein is conventionally described as a state in which the far-UV CD (which shows the secondary structure of a protein) is retained but the near-UV CD (which shows the tertiary structure of a protein) is lost. Dyes such as 8-anilino-naphthalene-1-sulfonic acid (ANS) exhibit a blue-shifted and enhanced fluorescence on binding to the MG state of a protein (and neither to the folded or unfolded forms). We have studied a metalloprotein azurin (reported to exhibit MG state), in the metal-free and metal-bound forms to investigate the same. Additionally, we have also studied ANS binding to some other common proteins (lysozyme, myoglobin, ubiquitin, and SUMO1) at physiological and low pH. We find that the holo-azurin exhibits ANS binding at a low pH, but partially chemically-denatured holo-azurin does not bind to ANS. At low pH, apo-azurin binds to ANS, but the protein is in the unfolded state (verified by CD and fluorescence). A variety of acid-denatured proteins also showed ANS binding. This may be due to ion-pair interactions between positively charged residues and the sulphonate group of ANS. We conclude that azurin does not exhibit an MG state at acidic pH. It must be noted that ANS binding alone is insufficient to establish the MG state of a protein, and usage of other methods is recommended. We further investigate the CP azurins (cpF114 and cpN42) for the presence of MG intermediates. Circular permutants can modify the energy landscape of proteins as per our previous studies with azurin.^73,74^ Studying molten globules in azurin permutants reveals how sequence rearrangements impact protein energetics and folding pathways. We find that Zn^2+^-cpN42 and apo-cpF114 azurins hint at the occurrence of MG states. Zn^2+^-cpN42 and apo-cpF114 both exhibit higher Trp mobility and solvent accessibility of the buried Trp as compared to WT azurin. We anticipate that our studies will facilitate the careful design of CPs for various applications.

## Supporting information

SuppInfo_

## ASSOCIATED CONTENT

**SUPPORTING INFORMATION:** Supporting information contains additional data, figures, and tables.

### Accession Code

Azurin UniProtKB-P00282.

### Conflict of Interest

The authors declare no conflict of interest.

### Funding Sources

Department of Atomic Energy (DAE), India, under project no. 12-R&DTFR-5.10-0100.

## Acknowledgments

S.R.K.A. acknowledges the financial support of the Department of Atomic Energy (DAE), India, under project no. 12-R&DTFR-5.10-0100. Plasmid (pGK22 vector) incorporated with wild-type azurin gene was a kind gift from Prof. G.W. Canters (Leiden University) and Plasmid (pET22b vector) incorporated with cpF114 azurin gene^71^ was a kind gift from Prof. Yang Yu (Tianjin Institute of Industrial Biotechnology, Chinese Academy of Sciences, China). We thank Mr. Aditya Shrivastava and Ms. Simran Arora of our research group who provided ubiquitin and SUMO1 proteins, respectively. We thank Mrs. Geetanjali Dhotre for her help in recording and analyzing MALDI-ToF data.

## ABBREVIATIONS

ANS: 8-Anilino-Naphthalene-1-Sulfonic acid
Asn: Asparagine
BSA: Bovine Serum Albumin
CD: Circular Dichroism
CP: Circular Permutation
Cys: Cysteine
DHFR: DiHydroFolate Reductase
EDTA: Ethylene Diamine Tetra Acetic acid
GdnHCl: Guanidine Hydrochloride
Gly: Glycine
His: Histidine
IRF: instrument response function
MALDI-ToF: Matrix Assisted Laser Desorption Ionization-Time of Flight
Met: Methionine
LMCT: Ligand to Metal Charge Transfer
MG: Molten Globule
NATA: N-Acetyl TryptophanAmide
Ni-NTA: Nickel-Nitrilo TriAcetic acid
MR: Nuclear Magnetic Resonance
PDB ID: Protein Data Bank IDentifier
SEC: Size Exclusion Chromatography
SUMO2: Small Ubiquitin-related MOdifier 2
TCE: trichloro ethanol
TCSPC: Time-Correlated Single Photon Counting
Trp: Tryptophan
UV: Ultraviolet
WT: Wild-Type.

## Notes

### Competing Interest Statement

The authors have declared no competing interest.

