## Supplementary material for "Circular permutants of azurin exhibit molten globule intermediates not observed in WT azurin": SuppInfo_

**Author ORCIDs**

Debanjana Das: 0000-0002-3178-7526

Sri Rama Koti Ainavarapu: 0000-0002-1646-2731

**Figure S1.**

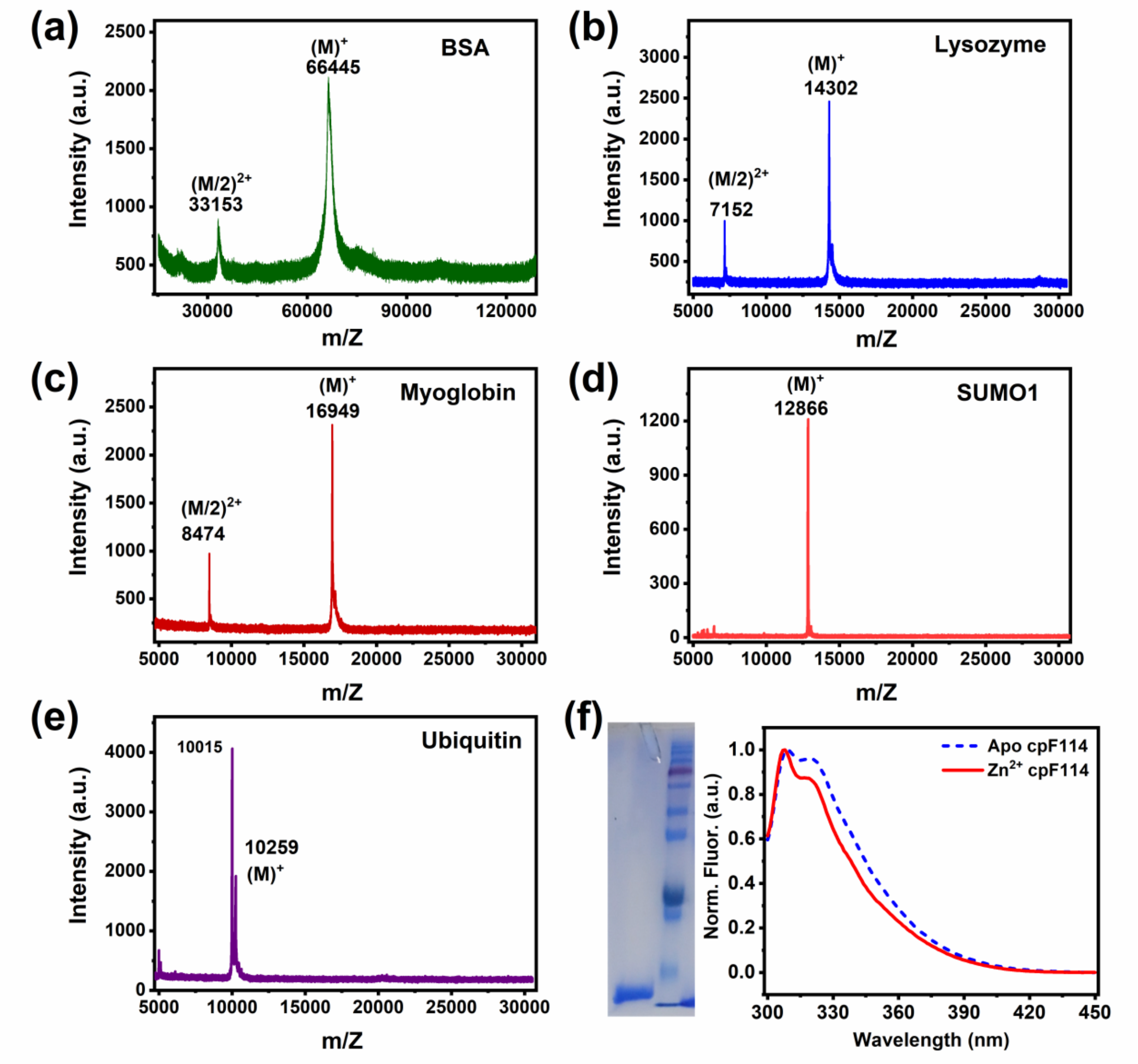


**Figure S1.** MALDI-ToF data for **(a)** BSA (bovine serum albumin), **(b)** lysozyme, **(c)** myoglobin, **(d)** SUMO1, and **(e)** ubiquitin, showing the molecular mass peak and $(M/2)^{2+}$ peaks. **(f)** (*Left*) Gel image of azurin-cpF114 in the first lane with the marker in the second lane (the lowest two bands of the marker are 11 and 17 kDa, respectively). (*Right*) Tryptophan fluorescence of apo- and $Zn^{2+}$-cpF114 azurin. Excitation of the samples was done at 295 nm.

#### Figure S2.

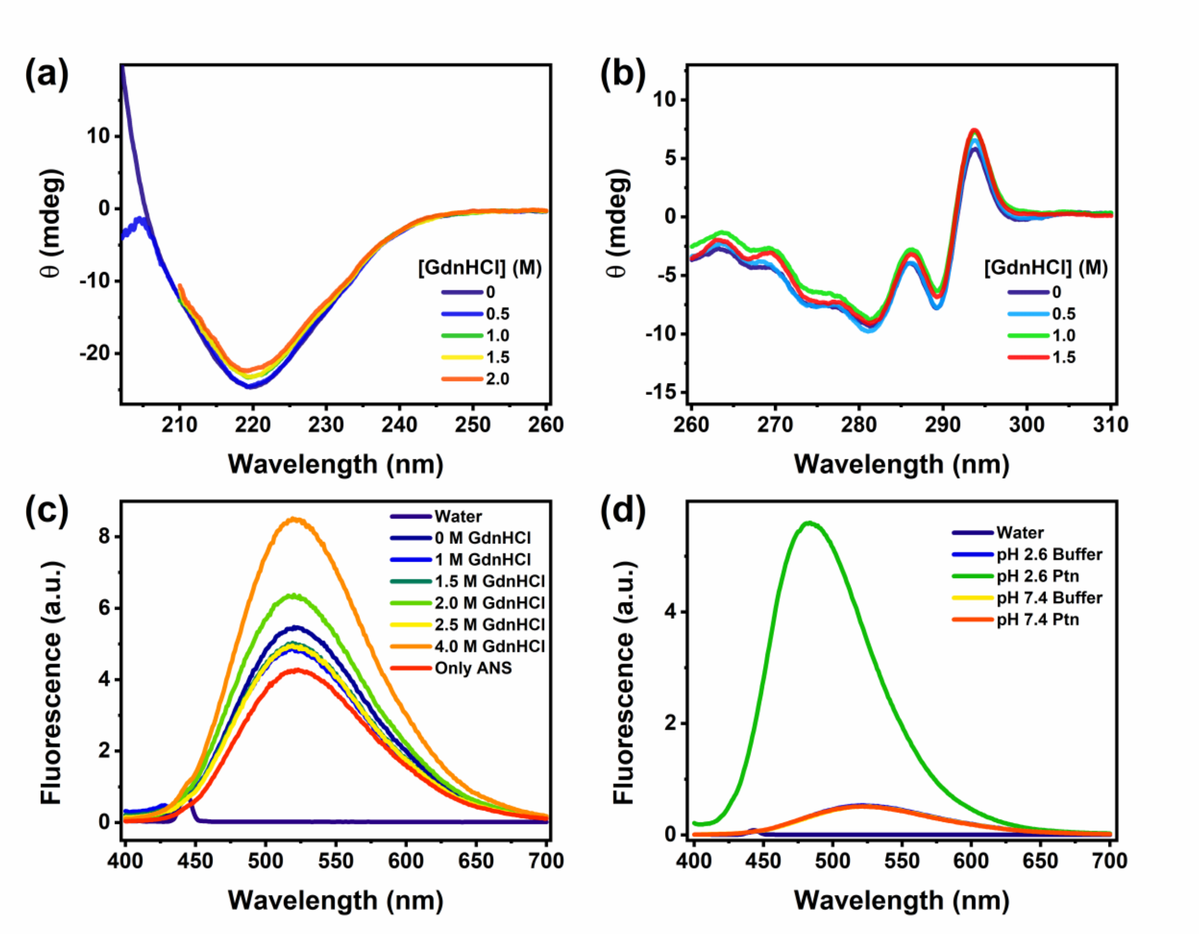


**Figure S2.** Chemical denaturation of $Zn^{2+}$-azurin WT probed by **(a)** far-UV CD, **(b)** near-UV CD, **(c)** steady state ANS fluorescence. **(d)** Steady-state fluorescence of ANS probing the acidic and neutral form of $Zn^{2+}$-azurin WT ([Protein] = 10 μM and [ANS] = 500 μM).

#### Figure S3.

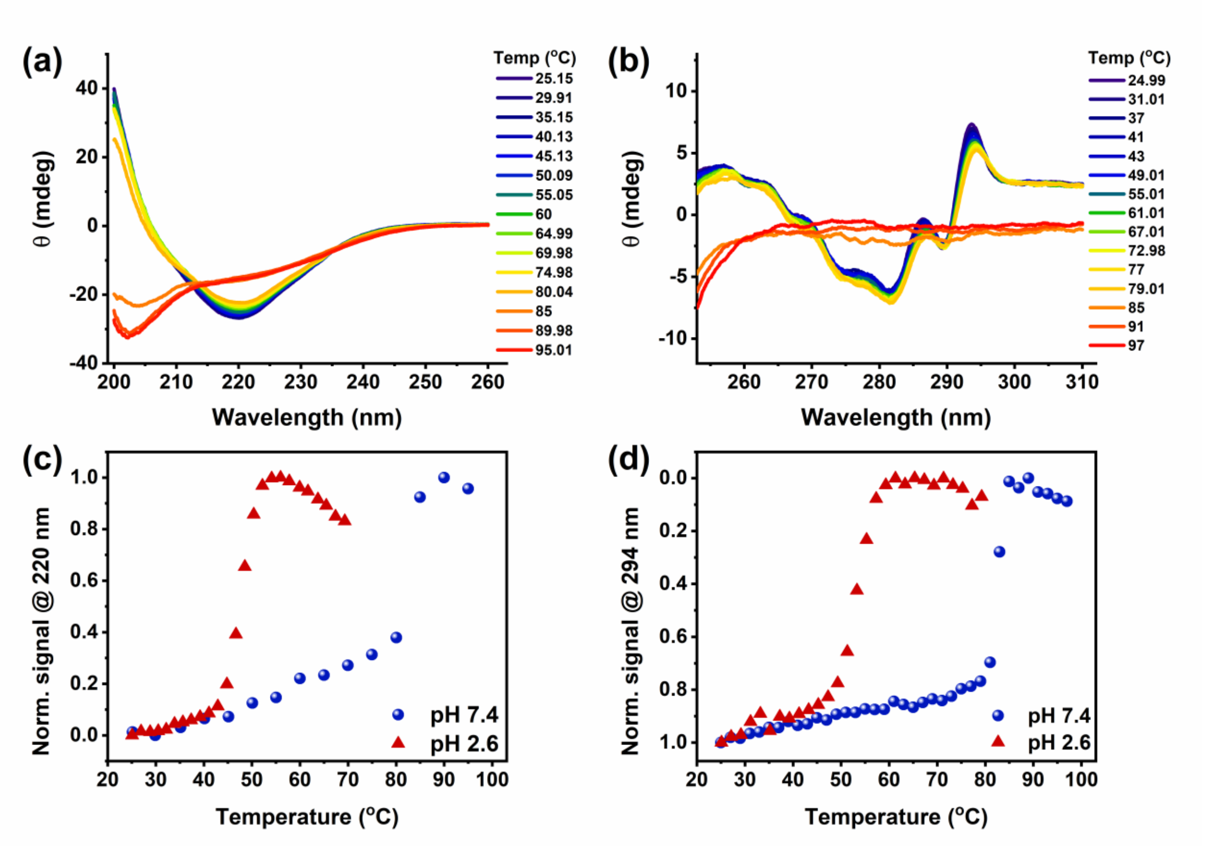


**Figure S3. (a)** Far-UV CD and **(b)** near-UV CD change of $Cu^{2+}$-azurin WT with temperature at pH 7.4. **(c)** Far-UV CD (at 220 nm) and **(d)** near-UV CD (at 294 nm) denaturation for $Cu^{2+}$-azurin WT at pH 7.4 (blue circles) and 2.6 (red triangles).

#### Figure S4.

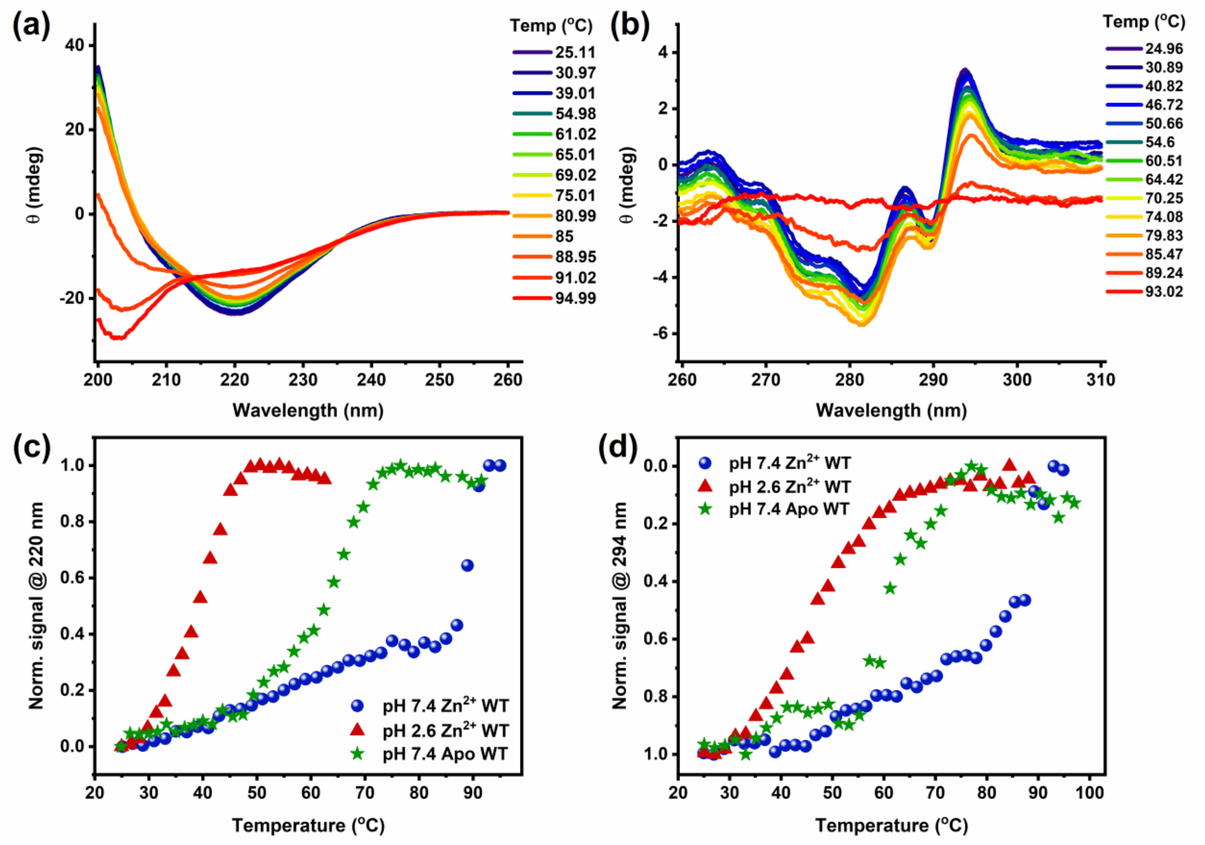


**Figure S4.** **(a)** Far-UV CD and **(b)** near-UV CD change of $Zn^{2+}$-azurin WT with temperature at pH 7.4. **(c)** Far-UV CD (at 220 nm) for $Zn^{2+}$-azurin WT at pH 7.4 (blue circles) and 2.6 (red triangles) and apo-azurin WT at pH 7.4 (green stars). **(d)** near-UV CD (at 294 nm) for $Zn^{2+}$-azurin WT at pH 7.4 (blue circles) and 2.6 (red triangles) and apo-azurin WT at pH 7.4 (green stars).

### Figure S5.

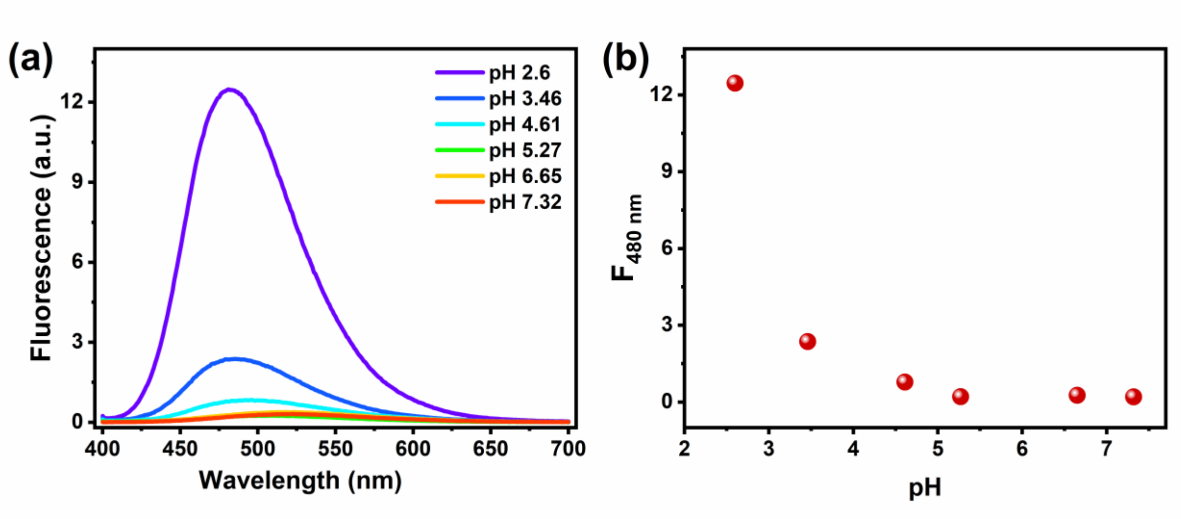


**Figure S5. (a)** ANS fluorescence response of apo-azurin WT with different pH solutions. **(b)** Fluorescence change of (a) at 480 nm with pH. ([Protein] = 10 μM and [ANS] = 500 μM).

### Figure S6.

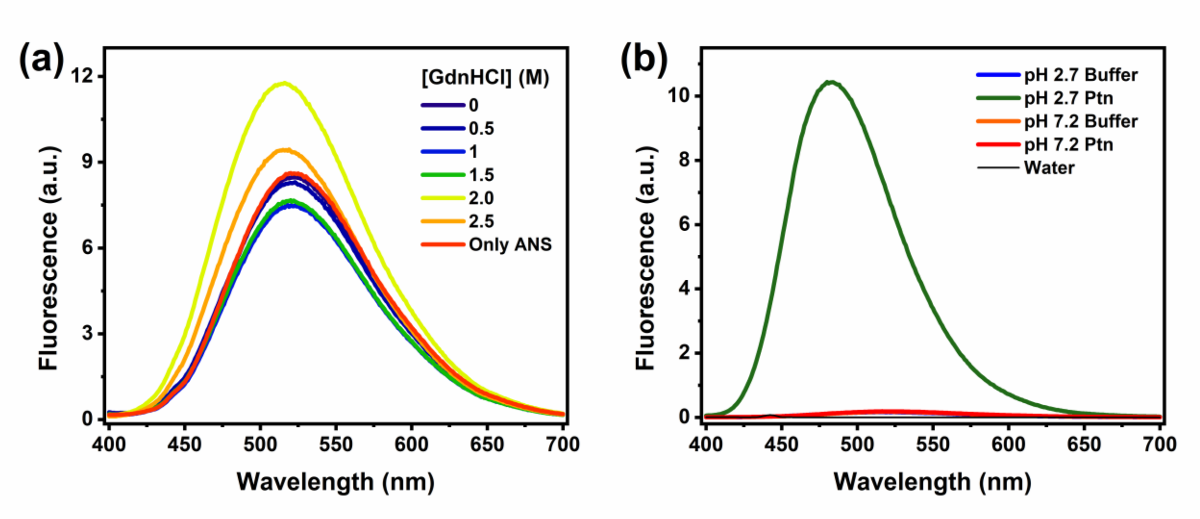


**Figure S6.** Chemical denaturation of $Cu^{2+}$-cpN42 azurin probed by steady-state fluorescence of ANS **(a)** on adding GdnHCl and **(b)** on reducing pH ([Protein] = 10 μM and [ANS] = 500 μM).

**Figure S7.**

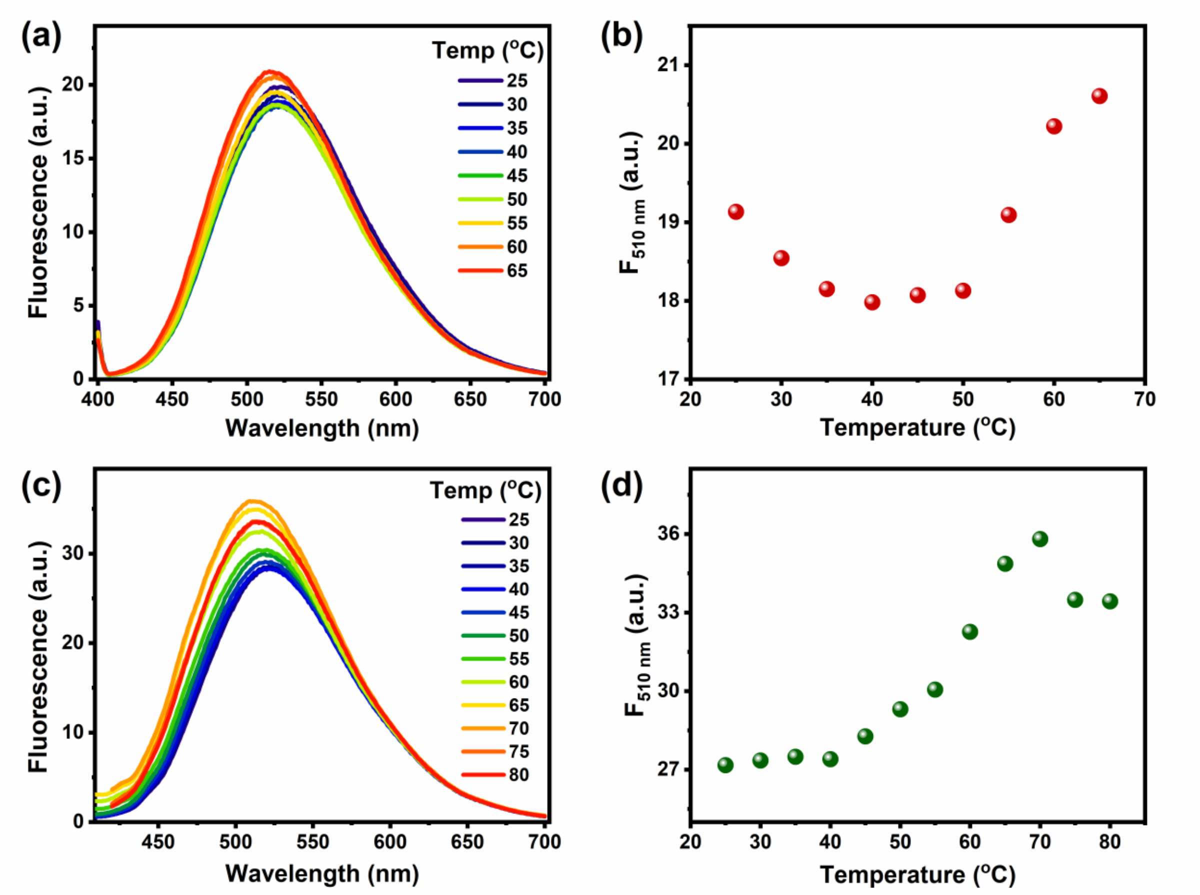


**Figure S7. (a)** ANS fluorescence response of apo-WT azurin with temperature change. **(b)** Fluorescence change of (a) at 510 nm plotted with temperature. **(c)** ANS fluorescence response of $Zn^{2+}$-cpF114 azurin with temperature change. **(d)** Fluorescence change of (c) at 510 nm plotted with temperature. [Protein] = 10 μM and [ANS] = 500 μM.

**Figure S8.**

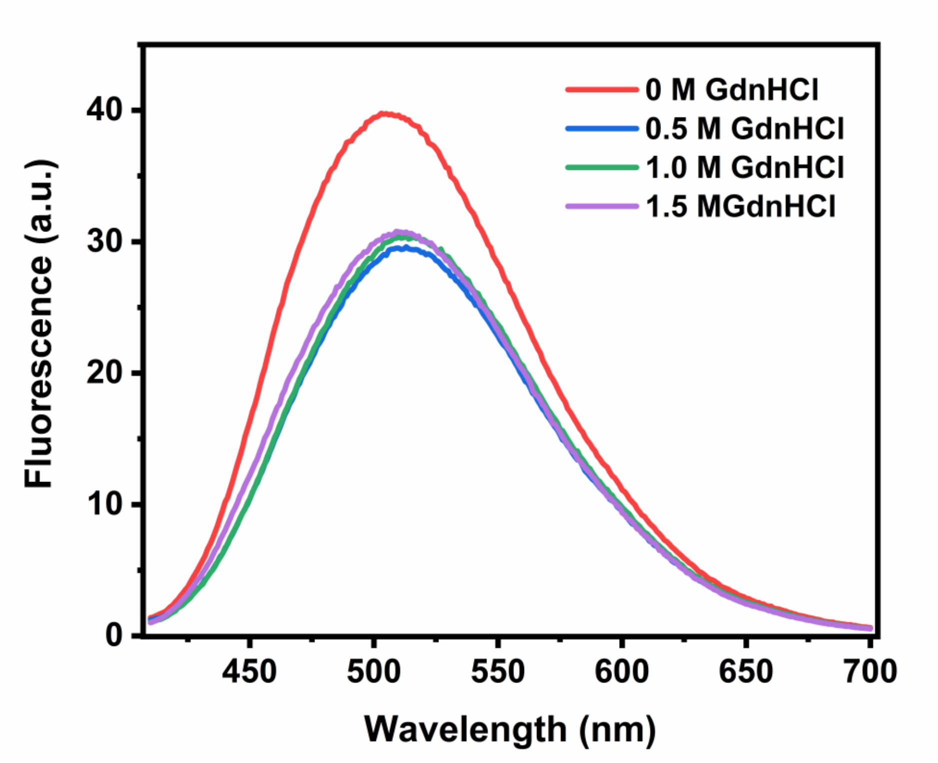


**Figure S8.** Chemical denaturation of $Zn^{2+}$-cpN42 azurin probed by steady-state fluorescence of ANS**.**

#### Figure S9.

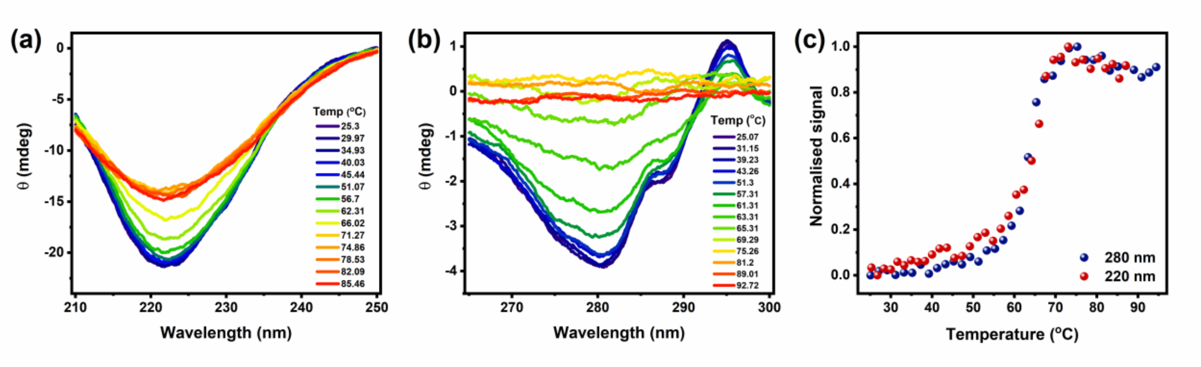


**Figure S9.** Change in **(a)** far-UV CD and **(b)** near-UV CD of $Zn^{2+}$-cpN42 azurin with temperature at pH 7.4. **(c)** Overlap of far-UV and near-UV CD of $Zn^{2+}$-cpN42.

**Figure S10.**

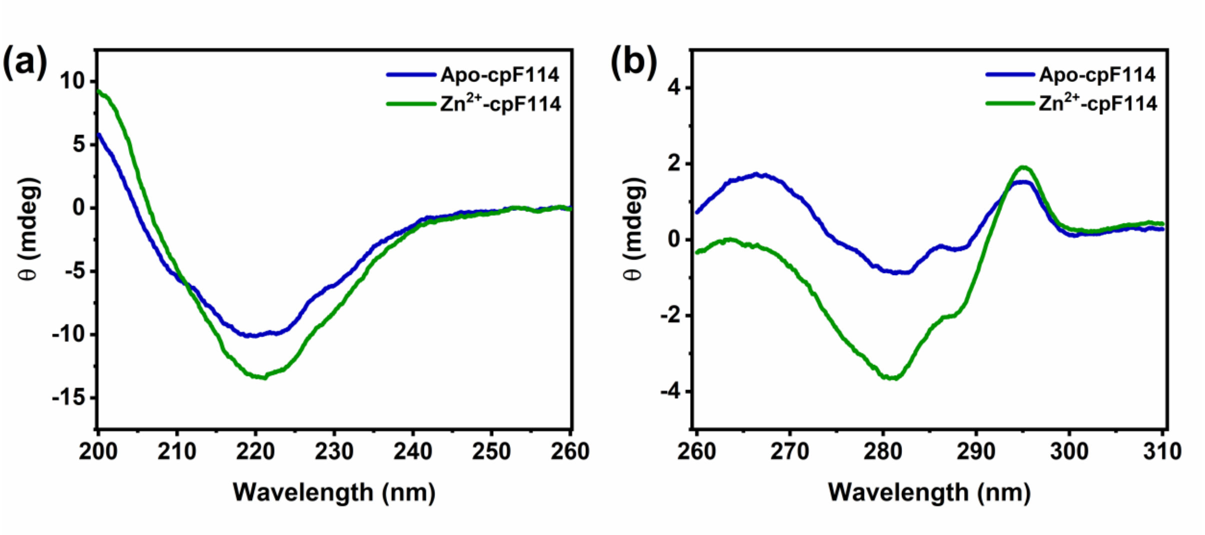


**Figure S10.** **(a)** Far-UV CD and **(b)** near-UV CD of apo-cpF114 and $Zn^{2+}$-cpF114 ($Zn^{2+}$ ion was added to the sample of apo-cpF114 and incubated for ~30 minutes before recording data).

#### Figure S11.

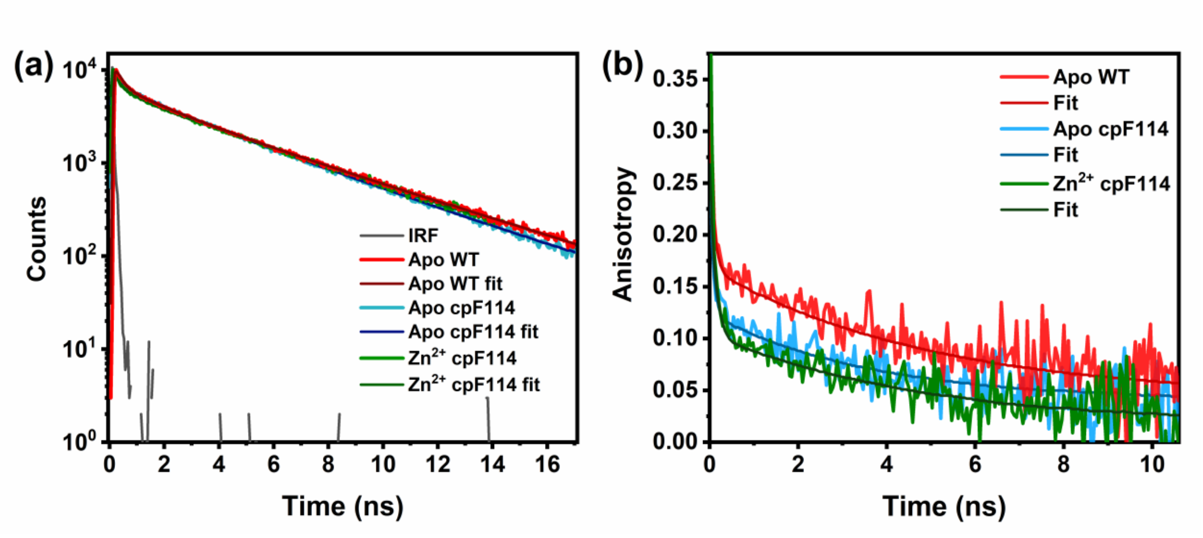


**Figure S11. (a)** Fluorescence lifetime data (with their corresponding fits) and **(b)** anisotropy decays (with their corresponding fits) of apo-cpF114, $Zn^{2+}$-cpF114, and apo-WT.

#### Figure S12.

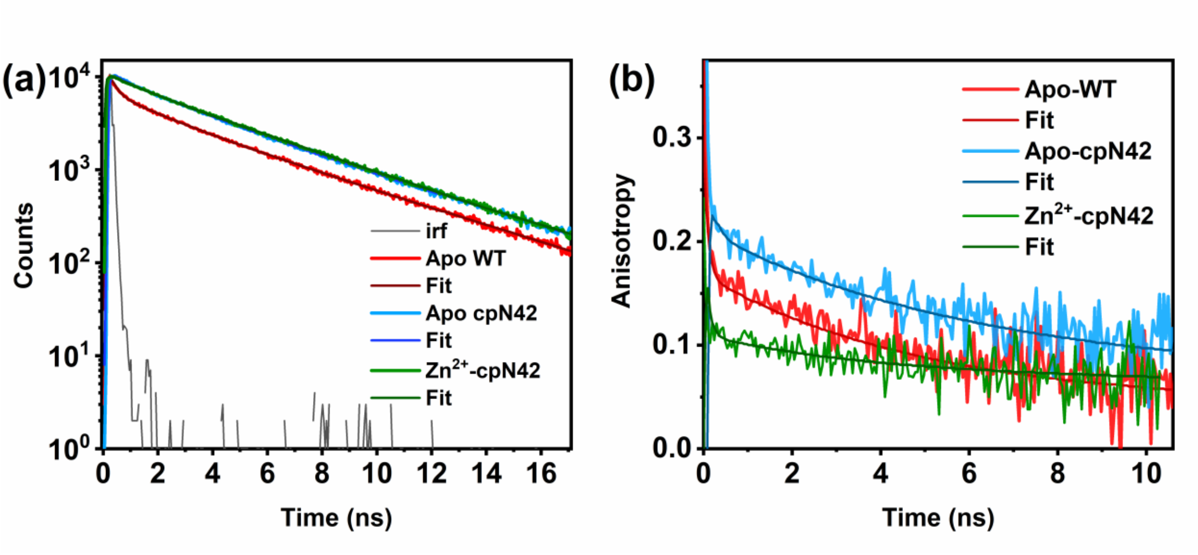


**Figure S12. (a)** Fluorescence lifetime data (with their corresponding fits) and **(b)** anisotropy decays (with their corresponding fits) of apo-cpN42, $Zn^{2+}$-cpN42, and apo-WT.

#### Supporting Information Tables

**Table S1.** Amino acid sequences of WT (derived from *Pseudomonas aeruginosa*), cpF114, and cpN42 azurins. Circular permutation (CP) proteins have linker regions highlighted in red.

| Protein | Amino acid sequences |
|---|---|
| Azurin WT | **AECSVDIQGN DQMQFNTNAI TVDKSCKQFT VNLSHPGNLP KNVMGHNWVL STAADMQGVV TDGMASGLDK DYLKPDDSRV IAHTKLIGSG EKDSVTFDVS KLKEGEQYMF FCTFPGHSAL MKGTLTLK** |
| Azurin cpF114 | **MGHSALMKGT LTLK**GIPGG**A ECSVDIQGND QMQFNTNAIT VDKSCKQFTV NLSHPGNLPK NVMGHNWVLS TAADMQGVVT DGMASGLDKD YLKPDDSRVI AHTKLIGSGE KDSVTFDVSK LKEGEQYMFF CTF** |
| Azurin cpN42 | MRGSHHHHHH **GSMGVMGHNW VLSTAADMQG VVTDGMASGL DKDYLKPDDS RVIAHTKLIG SGEKDSVTFD VSKLKEGEQY MFFCTFPGHS ALMKGTLTLK** GIPGG**AECSV DIQGNDQMQF NTNAITVDKS CKQFTVNLSH PGNLPK**RS |

**Table S2.** Far-UV structure analysis for apo-WT, apo-cpN42, apo-cpF114, $Zn^{2+}$-cpN42, and $Zn^{2+}$-cpF114 with the individual components in percentage using BeStSel analysis software.[1,2]

| Protein | α–helix | β–sheet antiparallel | β–sheet parallel | Turn | Others |
|---|---|---|---|---|---|
| Apo-WT | 3.5 | 31.8 | 4.1 | 10.8 | 49.8 |
| Apo-cpN42 | 2.5 | 35.3 | 2.1 | 13.5 | 46.7 |
| Apo-cpF114 | 1.3 | 37.3 | 0.0 | 16.2 | 45.2 |
| $Zn^{2+}$-cpN42 | 4.7 | 38.2 | 0.0 | 12.7 | 44.5 |
| $Zn^{2+}$-cpF114 | 2.6 | 39.0 | 0.0 | 15.6 | 42.8 |

**Table S3.** Fluorescence lifetime data of tryptophan of the azurin variants, collected at 340 nm wavelength (Tryptophan was excited at 295 nm).

| **Protein** | $\alpha_1$ | $\tau_1$ **(ns)** | $\alpha_2$ | $\tau_2$ **(ns)** | $\alpha_3$ | $\tau_3$ **(ns)** | $\tau_m$ **(ns)** |
|---|---|---|---|---|---|---|---|
| Apo-WT | 0.46±0.01 | 0.24±0.02 | 0.16±0.01 | 1.84±0.09 | 0.38±0.01 | 4.62±0.04 | 4.00±0.02 |
| Apo- cpF114 | 0.44±0.01 | 0.14±0.02 | 0.16±0.03 | 1.55±0.30 | 0.40±0.03 | 4.24±0.16 | 3.77±0.04 |
| $Zn^{2+}$- cpF114 | 0.52±0.02 | 0.11±0.01 | 0.12±0.01 | 0.72±0.14 | 0.36±0.02 | 4.20±0.04 | 3.88±0.02 |
| Apo-cpN42 | 0.23±0.11 | 1.30±0.66 | 0.56±0.16 | 3.81±0.95 | 0.41±0.18 | 5.00±0.32 | 4.12±0.03 |
| $Zn^{2+}$-cpN42 | 0.20±0.04 | 1.41±0.23 | 0.80±0.04 | 4.27±0.07 | - | - | 4.05±0.02 |

**Table S4.** Amplitudes of rotational correlation times of local motion ($\alpha_1$) and global motion ($\alpha_2$) of Trp in protein for WT and CP azurins. The amplitudes were used to calculate the semi-angle of rotation of Trp in the proteins.[3–5]

| Protein | $\alpha_1$ | $\alpha_2$ | Semi-angle of cone |
|---|---|---|---|
| Apo-WT | 0.54 ± 0.07 | 0.46 ± 0.07 | 39.9° |
| Apo-cpN42 | 0.48 ± 0.04 | 0.52 ± 0.04 | 36.8° |
| Apo-cpM13 | 0.78 ± 0.04 | 0.22 ± 0.04 | 53.8° |
| Apo-cpF114 | 0.65 ± 0.03 | 0.35 ± 0.03 | 45.8° |
| $Zn^{2+}$-cpF114 | 0.64 | 0.36 | 45.2° |
| $Zn^{2+}$-cpN42 | 0.81 ± 0.02 | 0.19 ± 0.02 | 56.0° |
